# Novel roles of small extracellular vesicles in regulating the quiescence and proliferation of neural stem cells

**DOI:** 10.1101/2021.08.20.456431

**Authors:** Jingtian Zhang, Junki Uchiyama, Koshi Imami, Yasushi Ishihama, Ryoichiro Kageyama, Taeko Kobayashi

## Abstract

Neural stem cells (NSCs) quiescence plays pivotal roles in securing sustainable neurogenesis and avoiding stemness exhaustion in the adult brain. The maintenance of quiescence and transition between proliferation and quiescence are complex processes associated with multiple niche signals, and environmental stimuli. Though the mechanisms of the transitions between NSC states have been extensively investigated, they remain to be fully elucidated. Exosomes are small extracellular vesicles (sEVs) containing functional units such as proteins, microRNAs, and mRNAs. It has already been demonstrated that sEVs actively participate in cancer cell proliferation and metastasis. However, the role of sEVs in NSC quiescence has not been investigated. Here, we applied proteomics to analyze the protein cargos of sEVs derived from proliferating, quiescent, and reactivating NSCs. Our findings revealed expression level fluctuations of NSCs sEV protein cargo at different proliferative conditions. We also identified functional clusters of gene ontology annotations from differentially expressed proteins in three sources of exosomes. Moreover, the use of exosome inhibitors revealed the contribution of exosomes to NSC quiescence at the entrance into quiescence, as well as in quiescence maintenance. Exosome inhibition delayed the entrance into quiescence by proliferating NSCs and allowed quiescent NSCs to exit from the G0 phase of the cell cycle. Protein translation was significantly upregulated in both quiescent NSCs and quiescent-induced NSCs via the exosome inhibition. Our results demonstrated that NSC exosomes are involved in regulating the quiescence of NSCs and provide a functional prediction of NSCs exosome protein cargos in terms of cell-cycle regulation and protein synthesis.

## 1 Introduction

NSCs, which are derived from the neuroepithelium of the neural tube, maintain the ability to self-renew and to give rise to neurons, astrocytes, and oligodendrocytes throughout a whole life. Two generally accepted canonical domains in the adult central nervous system that keep a reservoir of NSCs are the subventricular zone near the lateral ventricles and the subgranular zone of the dentate gyrus, where most of the NSCs remains quiescent (Cheung and Rando, 2013;Urban et al., 2019;Kobayashi and Kageyama, 2021). The quiescent state of NSCs, which is characterized by low metabolic rate and low protein and RNA synthesis, is vital to the preservation of their genomic integrity and essential functional properties (Cavallucci et al., 2016). Quiescent NSCs (qNSCs) rest at the G0 phase of the cell cycle and do not express proliferation markers, such as Ki-67 and MCM2 (Codega et al., 2014). However, they can be reactivated and re-enter the cell cycle upon proper signals. NSCs maintain a delicate balance with proliferating and quiescence. This equilibrium is essential for NSC homeostasis, and its disruption may lead to brain aging and its associated diseases (Cavallucci et al., 2016). Discrete NSC niche stimuli play indispensable roles in determining whether NSCs remain quiescent or reenter the cell cycle (Fuentealba et al., 2012;Kjell et al., 2020). Neurotrophin-3 is secreted by endothelial cells in the brain and choroid plexus and maintains NSC quiescence (Delgado et al., 2014), while Sphingosine-1-phosphate (S1P) and prostaglandin-D2 (PGD2) are G-protein coupled receptors in the cerebrospinal fluid (CSF) and actively sustain NSC quiescence (Codega et al., 2014). Though the transition between proliferating NSCs (active NSCs: aNSC) and qNSCs has been intensively studied, its mechanism has not been fully elucidated.

Extracellular vesicles are heterogeneous cell-derived membrane components that can be roughly categorized into small extracellular vesicles (sEVs, exosomes) and microvesicles (van Niel et al., 2018). sEVs are intraluminal vesicles (ILVs) formed by inward budding of the endosomal membrane during the maturation of multiple vesicle endosome (MVE), then are secreted outside of the cell by the fusion of plasma membrane and MVE. The sEVs vary widely in size but, like ILVs, are normally within the range of 30–100 nm and generally do not exceed 150 nm. Microvesicles are formed by direct budding from the plasma membrane and range in size from 100–1,000 nm (Colombo et al., 2014). Growing evidence has demonstrated the important roles of sEVs in signaling and regulating target cells via their diverse cargos, including protein, miRNA, and mRNA and various functional units (Valadi et al., 2007). The expression pattern of integrins in sEVs from cancer tissue differs in various kinds of cancers, which would partially account for the distinct metastatic destinies of various malignancies (Dolo et al., 1998; Hoshino et al., 2015). Ample evidence has proved that exosomes participate in the cell cycle and influence cell proliferation. Cancer cell-secreted exosomes increased the G1-phase cancer cell ratio and enhanced cell proliferation (Huang et al., 2019;Matsumoto et al., 2020). Exosomal miRNA, which is derived from the choroid plexus of the lateral ventricle, post-transcriptionally regulates quiescent NSC differentiation (Lepko et al., 2019). EVs secreted from NSCs have also been reported to be involved in Stat1 pathway in target cells and to act as a microglia morphogen (Cossetti et al., 2014;Morton et al., 2018). Moreover, EVs derived from differentiated neural stem/progenitor cells (NSPCs) have been reported to trigger the differentiation of proliferating NSPC (Stronati et al., 2019).

Despite the extensive research on the functions of EVs in NSCs (Batiz et al., 2015), the roles of sEVs in regulating the transition between aNSCs and qNSCs have not been certified. In the present study, we employed proteomics approaches to identify the functional differences among sEVs secreted from aNSCs, qNSCs and reactivating NSCs (reNSCs) *in vitro*. Our results demonstrated that the sEVs derived from the three sources differed in terms of protein content. From proteomics results, we hypothesized that NSCs might discard some proteins, such as ribosomes, to control the quiescence. To test this hypothesis, we treated NSCs with an inhibitor of sEV secretion during quiescence induction and resting. The inhibitor treatments facilitated NSC alterations from quiescence to proliferation. The expression of cell proliferation markers, MCM2, Ki-67, and CycD1, was upregulated in quiescent NSCs, as well as quiescence-induced NSCs, after inhibitor treatments. Exosome inhibition activated protein translation in quiescent and quiescent-induced NSCs, which might fit well with the “discarding model” in which some ribosomes are discarded to maintain low protein translation. Our findings confirmed that sEVs are involved in the maintenance of qNSCs, as well as their transition between proliferation and quiescence.

## 2 Methods

### 2.1 Cell culture, quiescence induction, and reactivation

NSCs were derived from ICR mouse embryos at E14.5 (Kobayashi et al., 2019). Active NSCs were cultured in proliferation medium [20 ng/mL epidermal growth factor (EGF); R&D systems, Inc., Minneapolis, MN, USA), 20 ng/mL bFGF (FUJIFILM Wako Pure Chemical Corporation, Osaka, Japan), penicillin/streptomycin (Nacalai Tesque, Kyoto, Japan), and N-2 Plus supplement (R&D Systems) in DMEM/F-12 (Gibco, Grand Island, NY, USA)] with 2 μg/mL laminin (Sigma-Aldrich Corp., St. Louis, MO, USA). Quiescence was induced with quiescent medium [50 ng/mL BMP4 (R&D Systems) in proliferation medium minus EGF] after two washings with phosphate-buffered saline (PBS; Nacalai Tesque) and achieved in 3 d. To prepare reNSCs, qNSCs were washed twice with PBS and cultured in proliferating medium for 2 d. For the exosome inhibition, aNSCs were cultured for 1 d and the medium was supplemented either with 10 μM GW4869 (Cayman Chemical Co., Ann Arbor, MI, USA) or dimethyl sulfoxide (DMSO; Nacalai Tesque) as the control. To measure protein synthesis, O-propargyl-puromycin (OP-puro, OPP; MedChemExpress Co., Monmouth Junction, NJ, USA) (Liu et al., 2012) was administrated for 1 h with 50 μM OP-puro in culture medium, followed by fixation with 4% paraformaldehyde (PFA; Nacalai Tesque) in PBS.

### 2.2 Exosome extraction

Supernatants from 15 cm culture dishes (Greiner, Oberosterreich, Austria) were centrifuged at 2,000 × g at 4°C for 20 min and at 10,000 × g at 4°C for 20 min and passed through a 0.22 μm filter (TPP, Trasadingen, Switzerland). The filtered supernatants were concentrated using a 100 kDa MW centrifugal filter (Amicon Ultra-15; EMD Millipore, Billerica, MA, USA) at 5,000 × g and then ultracentrifuged in a Beckmann optima XE with a SW 41 rotor (Beckman Coulter, Brea, CA, USA) at 100,000 × g at 4°C for 1 h. The pellets were washed with cold PBS and recentrifuged at 100,000 × g at 4°C for 1 h. The exosome pellets were recovered overnight in PBS at 4°C with gentle shaking and subjected to silver staining, western blot, particle analyses, and mass spectrometry. A total exosome isolation kit (Thermo Fisher Scientific, Waltham, MA, USA) was also used to collect exosomes from culture medium after inhibitor treatments by following the manufacturer’s protocol. The exosome pellets were recovered in PBS and subjected to NanoSight (Malvern Instruments, Malvern, UK) measurements. The exosome concentrations collected by the kit were 5.88E8+/- 6.06E6, 7.51E8+/-2.51E7, and 7.28E8+/-3.97E7 particles/mL from 2-d aNSC, 3-d qNSC, and 2-d reNSC cultures, respectively.

### 2.3 Protein quantification

Exosome proteins were quantified with a Micro BCA™ protein assay kit (Thermo Fisher Scientific). Bovine serum albumin (BSA) standard solutions and exosome samples were diluted in lysis buffer [50 mM Tris-HCl (pH 8.0), 100 mM NaCl, 5 mM MgCl2, and 0.5% (w/v) Nonidet P-40] and subjected to the manufacturer’s protocol. The protein concentrations in total cell lysates were quantified by the Lowry method using DC protein assay reagents (Bio Rad Laboratories, Hercules, CA, USA).

### 2.4 Western blot (WB) and silver staining

The cells were washed with cold PBS, lysed with lysis buffer [50 mM Tris-HCl (pH 8.0), 100 mM NaCl, 5 mM MgCl_2_, 0.5% (w/v) Nonidet P-40, Complete™ protease inhibitor cocktail (Roche Diagnostics, Basel, Switzerland), 1 mM phenylmethylsulfonyl fluoride, 250 U/mL Benzonase (Sigma), 10 mM β-glycerophosphate, 1 mM sodium orthovanadate, 1 mM NaF, and 1 mM sodium pyrophosphate] on ice for 30 min, and subjected to SDS-PAGE. The exosome samples were either directly mixed with SDS sample buffer or first precipitated using 10% (v/v) trichloroacetic acid (TCA) and resolved in sample buffer. The following primary antibodies were used for WB: rabbit anti-actin (Sigma-Aldrich), rabbit anti-CD9 (BioVision, Inc., Milpitas, CA, USA), rat anti-MFG-E8 (R&D Systems), mouse anti-CD63 (Novus Biologicals, Littleton, CO, USA), rabbit anti-Sox2 (EMD Millipore), and mouse anti-Cycd1 (Sigma-Aldrich). Horseradish peroxidase (HRP)-conjugated anti-rabbit, anti-mouse, and anti-rat (Thermo Fisher Scientific) were used as the secondary antibodies. Western blots (WBs) were visualized by chemiluminescence using Amersham ECL or ECL prime (Cytiva, Tokyo, Japan), quantified on an LAS3000 image analyzer (Fujifilm, Tokyo, Japan), and normalized against the corresponding intensity of β-actin for cell lysates. Statistical analyses were performed using GraphPad Prism 8 (GraphPad Software, La Jolla, CA, USA). For silver staining, a Pierce silver stain kit was used (Thermo Fisher Scientific).

### 2.5 Cell fixation and staining

Cells labeled with OP-puro were fixed with 4% PFA-PBS and stained with Alexa488-azide using Click-iT chemistry (Thermo Fisher Scientific). For immunocytochemistry to detect Ki-67 and MCM2, 4% PFA-fixed cells were immunostained with mouse anti-Ki-67 (BD Pharmingen, Franklin Lakes, NJ, USA) and rabbit anti-MCM2 (Abcam, Cambridge, UK) as the primary antibodies and then stained with anti-mouse-Alexa488 and anti-rabbit-Alexa594 as the secondary antibodies (Thermo Fisher Scientific) respectively and co-stained with DAPI (Sigma). Cell images were obtained on an AF6000 (Leica Microsystems GmbH, Wetzlar, Germany) and analyzed using ImageJ software (NIH, Bethesda, Maryland, USA). Statistical analyses were performed using GraphPad Prism 8 (GraphPad Software).

### 2.6 Exosome particle size measurement

Exosome particle size distribution and concentration were evaluated with NanoSight NS300 (Malvern Instruments). The size distribution was determined from five videos record by the NanoSight. For the exosome concentration determinations, ≤ 20 videos were recorded for more precise results (Parsons et al., 2017).

### 2.7 Transmission electron microscopy (TEM)

The exosome samples were subjected to negative staining by being loaded on the collodion surface of mesh grid (Nisshin EM, Tokyo, Japan), coated with 2% (w/v) uranyl acetate, and left to dry on the grid. They were then photographed under a JEM-1400 TEM (JEOL Ltd., Akishima-Shi, Japan).

## 3 Results

### 3.1 Purifications of exosomes secreted from aNSCs, qNSCs and reNSCs

We investigated the roles of sEVs secreted from NSCs in quiescent and proliferating conditions using cultured NSCs. To purify sEVs from cultured NSCs with high quality, we selected ultracentrifugation as the optimal method to collect abundant and pure sEVs (Fig. 1A). In our method, cell culture supernatants were centrifuged to remove cell debris and passed through a 0.22 μm filter to remove larger vesicles since sEVs is usually no larger than 150 nm in size. Nanoparticle tracking analysis (NTA) with NanoSight revealed that the estimated diameter of the purified sEVs was in the range of 100–130nm (Fig. 1B). Transmission electron microscopy (TEM) revealed their donut-shaped morphology (Fig. 1C). Results from both NTA and TEM analyses showed that the purified sEVs had typical exosomal characters. Previous reports used the term “exosome” to refer to a mixture of small vesicles because there was heretofore no method of fully separating them (Mathieu et al., 2019). In the present study, we use the term “exosome” to refer the sEVs. We then obtained exosomes from active, quiescent, and reactivating NSCs. BMP was used to induce quiescence in NSCs *in vitro* (Martynoga et al., 2013). Exosomes from aNSCs (aExo) were purified from aNSC culture supernatants after NSCs were cultured in dishes containing proliferating media for 2 d. Quiescence was induced by culturing NSCs in BMP-containing quiescent medium for 3 d and exosomes from qNSCs (qExo) were collected from the supernatants. QNSCs were reactivated by changing medium from quiescent to proliferating medium, and exosomes from reNSCs (reExo) were collected after the NSCs were cultured for 2 d. WB showed that exosome markers (CD9, CD63, and MFG-E8) (Thery et al., 2006) were expressed in the purified exosomes (Fig.1D). The transcriptional factor Sox2 was used as a negative control to rule out possible cell debris contamination (Fig.1D). Taken together, the foregoing data suggest that we successfully purified exosome samples suitable for use in the subsequent assays and experiments.

**Fig. 1.**
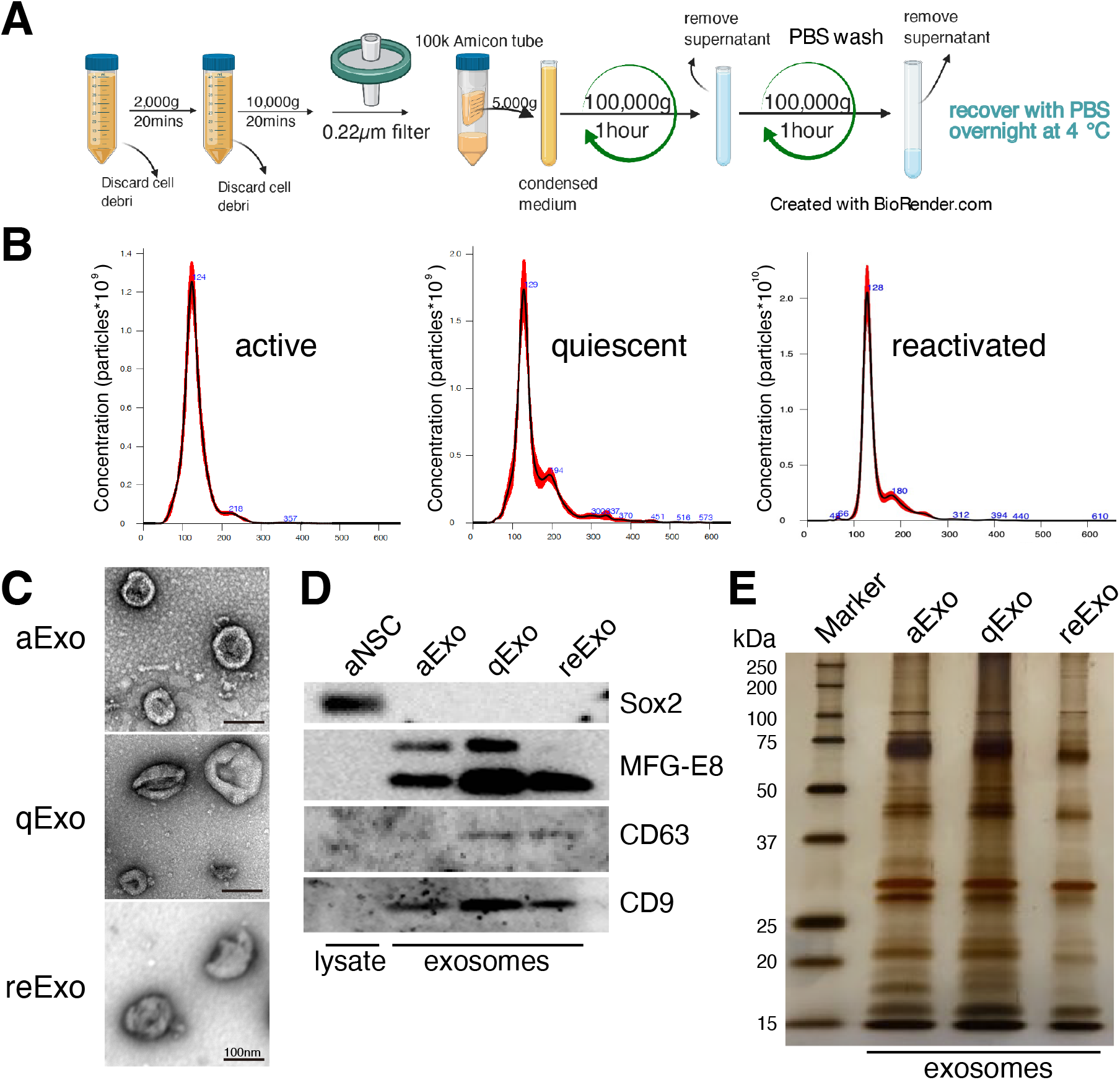
NSC exosomes collection and quality examination. (A) Exosome collection workflow. (B) Exosome size distribution and concentration. Exosomes were prepared from media for 2 d, 3 d, and 2 d incubation of active, quiescent, and reactivating NSCs, respectively. Averaged finite track length adjustment indicates exosome concentration and size, and was examined by NanoSight (n = 5). X-axis is vesicle size and y-axis is vesicle concentration. Numbers (blue) are approximate vesicles sizes at peak vesicle concentrations. (C) Transmission electron microscopy images of aExo, qExo, and reExo. Scale bar = 100 nm. (D) Total cell lysates and exosomes were subjected to WB of Sox2, MFG-E8, CD9, and CD63. MFG-E8 showing two bands indicating two variants. (E) Silver staining of exosome samples (0.4 μg). Left lane: molecular weight marker.

### 3.2 Proteomic profiling of exosomes derived from NSCs

We initially conducted silver staining to visualize proteomic inequalities and profiled the exosome protein contents. Silver staining showed similar protein expression patterns in the exosomes from active, quiescent, and reactivating NSCs (Fig. 1E). We then exanimated the exosome profiles by nano-scale liquid chromatography/tandem mass spectrometry (nano LC/MS/MS) (Supplementary methods). Digested proteins from individual exosome samples were labeled with nine different Tandem mass tag (TMT) 10-plex reagents (aExo vs. qExo vs. reExo; n = 3 each) (Ogata and Ishihama, 2020). Relative protein abundance was quantified based on the “reporter ion intensity.” Among 1,283 proteins we identified, 1,178 proteins were quantified in at least two out of the three replicates in at least one condition and were used for further analysis (Supplementary Table 1). To compare the exosome contents in the databases, we consulted the lists of proteins in Exocarta which includes those involved in MVB biogenesis, those associated with membrane proteins such as CD9 and CD63, exosome markers, and so on (Keerthikumar et al., 2016). There were 372 proteins common to the NSC exosomes and Exocarta (Bardou et al., 2014). Thus, our exosome samples shared many proteins with those previously reported (Fig. 2A). The sample comparisons were “aExo vs. qExo” (AQ), “qExo vs. reExo” (QRE), and “aExo vs. reExo” (ARE) (Fig. 2B and 2C). The volcano plots revealed bilateral significantly expressed proteins (log2 fold change (FC) >1; p-value < 0.05) in AQ and QRE (Fig. 2B; left and center charts). For ARE, however, there were only eight types of highly enriched proteins in aExo and no upregulated proteins in reExo relative to aExo (orange dots in right chart of Fig. 2B; Supplementary Table 2).

**Fig. 2.**
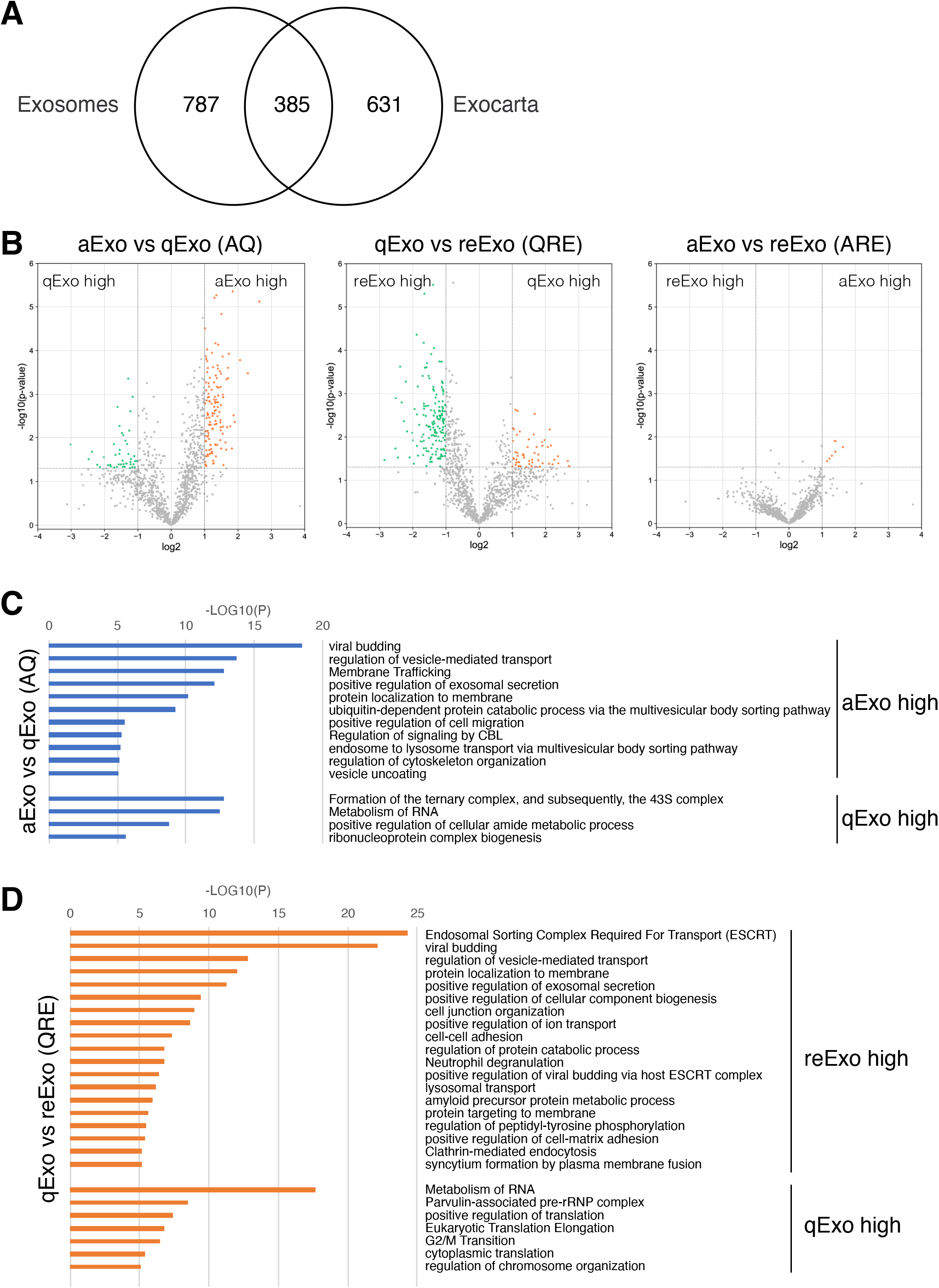
Exosome proteomes and gene ontology (GO). (A) Venn diagram of proteins identified in exosomes derived from all NSCs and the exosome proteome database Exocarta. (B) Volcano charts of differentially expressed exosomal proteins in aExo vs. qExo (AQ, left chart), qExo vs. reExo (QRE, center chart), and aExo vs. reExo (ARE, right chart). Orange and Green dots: significantly enriched proteins. (C, D) GO analyses of significantly enriched proteins indicated by green and orange dots in panel B.

To obtain functional insights of the differentially expressed protein cargos, we used gene lists of each comparison groups (log_2_ FC > 1; p-value < 0.05) in a gene ontology (GO) analysis (Supplementary Table 2) conducted on the Metascape website (https://metascape.org/gp/index.html#/main/step1). We identified enriched GO/KEGG and Reactome terms (Zhou et al., 2019). For both AQ and QRE, vesicle trafficking-related terms were significant in aExo, and translation-related terms were significant in qExo (Fig. 2C; Supplementary Table 3). No GO analysis was conducted on ARE as there were too few enriched factors, and the functional clusters could not be calculated. We also applied the “molecular complex detection” (MCODE) algorithm to detect densely related regions in the pool of protein-protein interaction webs (Bader and Hogue, 2003) and explore potential connections among our exosome proteomes. The algorithm speculated high-confidence networks of highly enriched proteins in AQ (Fig. 3A and 3B) and QRE (Fig. 3C and3D) and applied the enriched terms to annotate each MCODE network (Supplementary Tables 4). MCODE cluster identified endocytosis- and membrane trafficking-related clusters as aExo-enriched terms and translation- and ribosome-related clusters as the qExo-enriched terms in both AQ and QRE comparisons. It was consistent with the elevated scores of ribosomal protein subunits in qExo proteome (Supplementary Table 4). The foregoing results disclosed the contrasts between the quiescence and proliferation stages and facilitated the exploration of potential functional units from exosome proteomes.

**Fig. 3.**
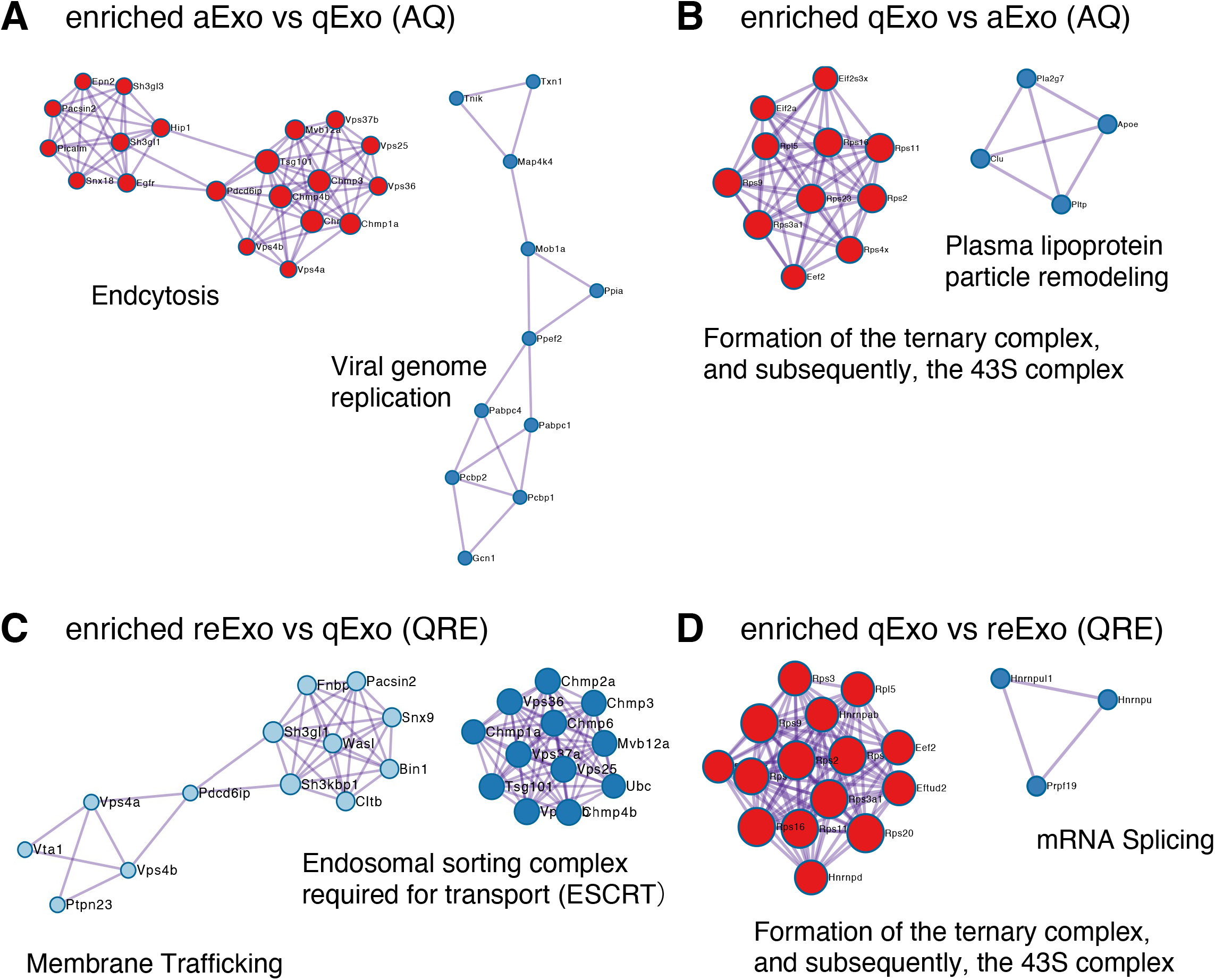
Molecular complex detection networks among enriched proteins. Molecular complex detection networks of putative protein-protein interactions among enriched proteins in exosomes from aNSCs (aExo), exosomes from qNSCs (qExo), and exosomes from reNSCs (reExo). Each node color indicates a different network cluster. (A) Annotations enriched in aExo in aExo vs. qExo (AQ). (B) Annotations enriched in qExo in AQ. (C) Annotations enriched in reExo in qExo vs. reExo (QRE). (D) Annotations enriched in qExo in QRE.

### 3.3 Exosomes participate in quiescence regulation of NSCs

Our results demonstrated distinct differences in exosome protein contents and enriched pathways between the quiescence and proliferation stages of NSCs. We speculated that the high enrichment of translation-related proteins in qExo may imply the contribution of exosomes to reduce protein translation in qNSCs by discarding the contents from cells. To explore the role of exosomes in this hypothesis, we applied a chemical inhibitor of the exosome (Fig. 4A). We used GW4869, a neutral, noncompetitive inhibitor of sphingomyelinase (N-SMase) which reduces ceramide levels and inhibits the ceramide-mediated release of mature exosomes from MVBs (Verderio et al., 2018). GW4869 is reported to inhibit exosome secretion by HEK293 cells, macrophages, and mesenchymal stem cells (Kosaka et al., 2010;Essandoh et al., 2015;Che et al., 2019). Nanosight measurements revealed that approximately one third of the exosome secretion was blocked in NSCs by GW4869 (Fig. 4B). We selected GW4869 for our subsequent analyses and investigated its effects.

**Fig. 4.**
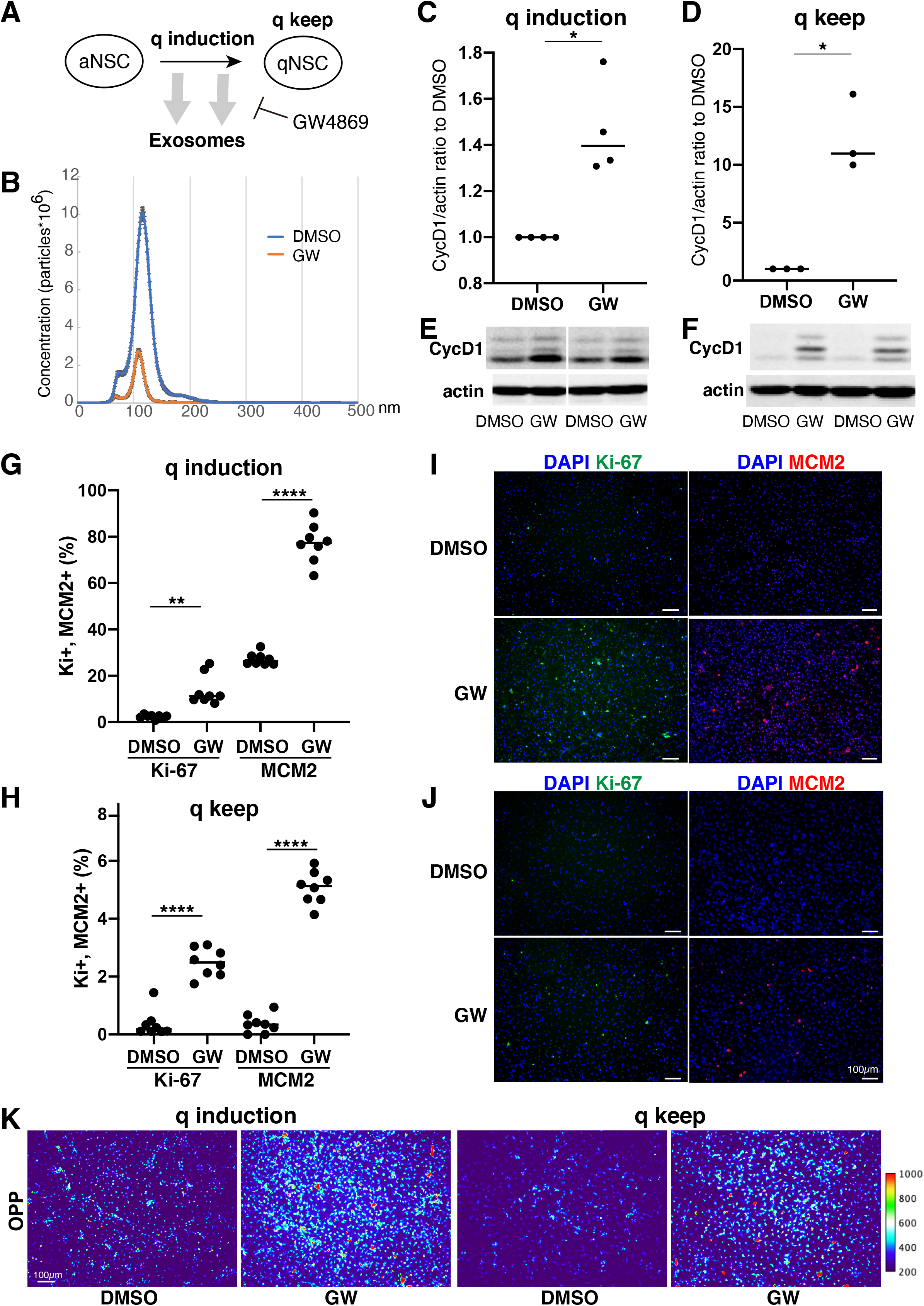
Exosomes involved in cell cycle regulation of qNSCs. (A) Exosomal functions in quiescence entrance and maintenance. (B) Exosome quantification of aNSCs after treatment with GW4869 (orange) and DMSO as a control (blue). Horizontal axis: vesicle size. Vertical axis: vesicle number. (C, D) Quantification of CycD1 levels in NSCs after inducing (C) and maintaining quiescence (D) in presence of GW4869 for 12 h (C) and 24 h (D). All values were divided by that for DMSO control. Each plot indicates independent experiments. CycD1 band intensities were divided by those of actin bands. Ratios with respect to DMSO control are plotted. Middle lines in plots are medians. (\**P* < 0.05, *\*\*P* < 0.01; two-tailed paired t-test, n = 4; For q-keep, one outlier identified by outlier test of Grubbs was removed.) (E, F) Representative results of WB of quiescence induction (E) and maintenance (F). CycD1 were three bands, and actin is loading control. (G, H) Cell counts of Ki-67- and MCM2-positive cells after inducing (G) and maintaining quiescence (H) after incubation with GW4869 for 24 h. Ki-67- and MCM2-positive cell numbers were divided by the cell number based on DAPI staining and shown as the percentage. (\*\**P =* 0.013, *\*\*\*\*P <* 0.0001; one-way ANOVA, Turkey’s multiple comparison test, n = 8). (I, J) Representative photos of Ki-67- (green) and MCM2-positive cells (red) with DAPI staining (blue) in panel G (I) and H (J). (K) Protein synthesis activity in the quiescence-induced (q induction) and maintained cells (q keep). The intensity of OP-puro labeling (OPP) is represented by a thermal scale.

We next investigated whether reduced exosome secretion affects the induction of NSC quiescence in proliferating NSCs and the maintenance of quiescence in qNSCs. We tested CycD1 expression levels by WB analysis for the NSC proliferation criteria (Baldin et al., 1993). We treated NSCs with GW4869 and DMSO, as a negative control, dissolved in the quiescent medium. Contrasting CycD1 expression was detected 12 h after inducing quiescence with the exosome inhibitor. The cells treated with GW4869 displayed higher CycD1 expression levels than those in the DMSO control (Fig. 4C and 4E). Hence, interference of exosome biogenesis and secretion impeded the NSCs from entering the G0 phase. We treated qNSCs with GW4869 in the quiescent medium to certify the influence of hindered exosome activity on the maintenance of quiescence. We discovered that after 24 h, CycD1 expression was upregulated in the presence of GW4869 (Fig. 4D and 4F). We investigated two proliferation markers, Ki-67 and MCM2, which are widely used to detect proliferating NSCs, in GW4869-treated NSCs. MCM2 is highly expressed throughout the cell cycle, including the G1 phase and except the G0 phase, whereas Ki-67 expression peaks at M phases and is low at the G1 phase (Miller et al., 2018;Harris et al., 2021). Exosome inhibition increased the percentages of proliferating (Ki-67+ and MCM2+) cells in both quiescent-induced and quiescent-mainteined NSCs (Fig. 4G–J). Thus, the blockage of exosome secretion prevented the entry of proliferating NSCs into the G0 phases and destablized the G0 phase.

To elucidate the mechanism through which exosomal inhibition destabilized the G0 phase in NSC quiescence, protein translation was measured using a puromycin analogue, OP-puro (OPP). OPP is effectively incorporated into newly generated proteins and can be used to quantify the protein synthesis rates in cells by detecting OPP-labeled proteins after fixation (Liu et al., 2012). aNSCs and qNSCs were incubated in GW-containing quiescent medium for 24 h, labeled with OPP for 1 h, and analyzed. Exosome inhibition increased the levels of OPP-labeled proteins in quiescent-induced and quiescent-maintained NSCs compared to those in the controls (Fig. 4K). This higher translation in GW-treated NSCs was consistent with higher percentages of proliferating cells in those than in controls under quiescent culture conditions (Fig. 4G–J). Taken together, these results demonstrated a strong correlation between exosome secretion and cell cycle regulation together with protein synthesis in NSCs quiescence.

## 4 Discussion

Exosomes are ubiquitously secreted by nearly all cell types and thought to actively participate in various niche signal transductions (Thery et al., 2009;Rashed et al., 2017;Bonafina et al., 2020). In this study, we successfully purified exosomes from NSCs that were available for proteomics (Fig. 1). The exosomal proteomes we acquired contained not only many common proteins with those previously reported but also uncommon proteins that may specifically exist in NSCs (Fig. 2). Moreover, our results highlighted the different cargo in exosomes from the same NSC origin but at different states (Fig. 3). Exosomes are known as biomarkers of cancers to inform the origins and biological changes. Our MS data might contribute to specify the condition and maintenance of NSC activation and quiescence in the adult brain in future.

We initially tested several exosome collection kits and methods but discovered that they introduced protein contamination. Ultracentrifugation provided high-purity exosomes but the yield was too low due to sample loss during the procedure (Fig. 1). Hence, we increased the cell number, culture medium volume, and recovering time after ultracentrifuge and finally obtained high-purity, high-yield exosomes. We then conducted TMT-based quantitative proteomics by nanoLC/MS/MS analyses to clarify the roles of exosomes from NSCs. A GO analysis of the enriched proteins at each state disclosed functional clusters that facilitated determination of the roles of exosomes (Fig. 2 and 3). It was established that vesicular transport and ribosomes were enriched pathways in the proliferating and quiescent exosomes, respectively.

A prevalence of ribosomal proteins has been reported for sEVs from cancer cells (Mathivanan et al., 2010;Ji et al., 2013;Willms et al., 2016). Our study showed that there was apparent concentrated expression of 40S ribosomal subunit, which forms a pre-initiation complex for translation (Jackson et al., 2010) in qExo. As quiescent cells have low protein synthesis rates, exosomes may export translation-related components from cells during quiescence induction. The “discarding model” of exosome function upon entering quiescence may explain the fact that ribosomal protein expression levels rise in exosomes but fall in cells (Pan et al., 1985). However, the discarding model does not rule out the possibility that the exosome cargos may transfer signals to recipient cells or alter their microenvironments. Future research should explore the molecular function of specific proteins within our exosome proteomes as some of them might be essential for tuning cell proliferation from quiescence. For example, PCSK6 (also called PACE4) (Supplementary Table 2), enriched in aExo, is a protein convertase that could promote cell proliferation by inducing the ERK1/2 and STAT3 signaling pathways (Seidah et al., 2013;Jiang et al., 2017). It would be an attractive target if the specific factor or exosomes may reactivate NSCs in the adult brain in order to avoid the functional decline of brain with aging and brain disorders.

Finally, our study revealed that exosome production and secretion regulate NSC quiescence using the exosome inhibitor GW4869 (Fig. 4). First, we found that the exosome inhibitor increased CycD1 levels in qNSCs, which indicated the re-entry of qNSCs to the cell cycle, and prevented the reduction in CycD1 in aNSCs during the entrance to the G0 phase after quiescence induction. Second, the exosome inhibitor significantly increased the number of proliferating cells expressing Ki-67 and MCM2, even under quiescent culture conditions. Third, protein translation was greatly upregulated in qNSCs and quiescence-induced NSCs. These results suggested that interference with exosome biogenesis and secretion impedes the maintenance of and transition to quiescence in NSCs by upregulating translation. The “discarding model” we described previously herein might be appropriate based on the results indicating that exosomes exhaust the functional translational machinery to reduce new protein synthesis for proliferation. Further research will be required to confirm this hypothesis and to identify the molecular mechanisms connecting proliferation with protein synthesis. A recent report demonstrated that proliferating NSCs gradually enter shallow quiescence with age and transfer between two states, quiescence and proliferation, and the balance is regulated by the degradation of a specific transcriptional factor, Ascl1 (Harris et al., 2021). Exosomal regulation of new protein synthesis, we hypothesized here, is potentially a novel mechanism regulating this balance in the adult brain. Taken together, our results established the functional importance of exosomes in participating in the quiescence regulation of NSC quiescence.

## Author contributions

JZ and TK designed and conducted all experiments. JU, KI, and YI performed the proteomic analyses. JZ, TK, KI, and RK wrote the manuscript.

## Abbreviations

aExo: aNSC exosomes
aNSCs: active neural stem cells
AQ: aExo vs. qExo
ARE: aExo vs. reExo
BMP: bone morphogenic protein
CSF: cerebrospinal fluid
DMSO: dimethyl sulfoxide
EGF: epidermal growth factor
ESCRT: endosomal sorting complexes for transport
GO: gene ontology
KEGG: Kyoto Encyclopedia of Genes and Genomes
HRP: horseradish peroxidase
ILVs: intraluminal vesicles
LC: liquid chromatography
MCODE: molecular complex detection
miRNAs: microRNAs
MS: mass spectrometry
NSPCs: neural stem/progenitor cells
MTA: nanoparticle tracking analysis
MVE: multiple vesicle endosome
nano LC/MS/MS: nanoscale liquid chromatography-tandem mass spectrometry
NSCs: neural stem cells
N-Smase: neutral-sphingomyelinase
OPP: O-propargyl-puromycin (OP-puro)
PGD2: prostaglandin-D2
qExo: qNSC exosomes
qNSCs: quiescent neural stem cells
QRE: qExo vs. reExo
reExo: reNSC exoxomes
reNSCs: reactivating neural stem cells
sEVs: small extracellular vesicles
S1P: sphingosine-1-phosphate
SDS-PAGE: sodium dodecyl sulfate polyacrylamide gel electrophoresis
TCA: trichloroacetic acid
TEM: transmission electron microscopy
TMT: tandem mass tag
WB: western blotting

## Acknowledgements

The authors thank P. Kurre, J. Hejna, Y. Okuno, H. Kouda, K. Furuta-Okamoto, K Ishii and members of the Kageyama Laboratory for their technical assistance and discussions.

## Data availability

MS raw data and analysis files have been deposited to the ProteomeXchange Consortium (http://proteomecentral.proteomexchange.org) via the jPOST partner repository (https://jpostdb.org) (Moriya et al., 2019) under the data set identifier PXD027651. Login information to jPOST: https://repository.jpostdb.org/preview/9491816326103a2b48aec3, access code 7870.

## Supplementary Materials

### Supplementary Methods

**Supplementary Table 1. Proteins identified using proteomic analysis.**

**Supplementary Table 2. Differential expression of proteins in each exosome sample.**

Intensity 1–3 represents triplicate samples of exosomes from aNSCs (aExo samples). Intensity 4–6 represents triplicate samples of exosomes from qNSCs (qExo samples). Intensity 7–9 represents triplicate samples of exosomes from reNSCs (reExo samples).

**Supplementary Table 3. Gene ontology analysis of enriched proteomes.**

**Supplementary Table 4. Gene ontology annotation lists.**

