## Supplementary Methods for "Novel roles of small extracellular vesicles in regulating the quiescence and proliferation of neural stem cells"

### 1. Sample pretreatment for mass spectrometry (MS)

Protein samples (1.5 µg each) were dried with a SpeedVac (Thermo Fisher Scientific) and resuspended in a phase transfer surfactant buffer containing 12 mM sodium deoxycholate, 12 mM sodium *N*-lauroylsarcosinate, 100 mM Tris-HCl (pH 9.0) and protease inhibitor cocktail (Merck KGaA, Darmstadt, Germany) (Masuda et al., 2008). The proteins were reduced with 10 mM dithiothreitol at 37°C for 30 min and alkylated with 50 mM of 2-iodoacetamide at 37°C in the dark for 30 min. After fivefold dilution in 50 mM ammonium bicarbonate, the proteins were digested by 0.1 µg lysyl endopeptidase (Lys-C) and 0.1 µg trypsin (Promega, Madison, WI, USA) overnight at 37°C. The samples were acidified with 0.5% (v/v) trifluoroacetic acid (TFA) and desalted using StageTips packed with SDB-XC Empore disk membranes (GL Sciences, Tokyo) (Rappsilber et al., 2003). All chemicals used here were purchased from FUJIFILM Wako Pure Chemical Corporation.

### 2. Tandem mass tag (TMT) labeling and high-pH reversed-phase LC fractionation

The desalted peptides were evaporated and labeled with 10-plex TMTs (Thermo Fisher Scientific) as previously described (Ogata and Ishihama, 2020). The TMT-labeled peptides were separated into eight fractions by high-pH, reversed-phase LC fractionation usign SDB-XC StageTips. The fractionated peptides were dried and resuspended in a loading solution (0.5% (v/v) TFA and 4% (v/v) acetonitrile (ACN)) for subsequent nano-scale liquid chromatography/tandem MS (nanoLC/MS/MS) analyses.

### 3. NanoLC/MS/MS analyses

The peptides were analyzed with an Orbitrap Fusion Lumos mass spectrometer (Thermo Fisher Scientific), coupled with a Thermo Ultimate 3000 RSLCnano pump, an HTC-PAL autosampler (CTC Analytics, Zwingen, Switzerland) and a self-pulled analytical column (150 mm L × 100 µm i.d.) packed with ReproSil-Pur C18-AQ (3 µm; Dr. Maisch GmbH, Ammerbuch, Germany) (Ishihama et al., 2002). The injection volume was 5 µL and the flow rate was 500 nL/min. The mobile phases consisted of (A) 0.5 % (v/v) acetic acid and (B) 0.5 % (v/v) acetic acid and 80 % (v/v) ACN. The gradient was as follows: 5–15 % B in 5 min, 15–40 % B in 60 min, 40-99 % B in 5 min, and 99 % B in 5 min. The splay voltage was 2.4 kV. The MS was operated in data-dependent acquisition with a full scan in the Orbitrap followed by 3-s MS/MS scans. Full scans were performed from *m/z* 375–1,500 with 120,000 resolutions after accumulation to a 4 x $10^5$-ion target value and 50 ms maximum injection time. The MS/MS scans were performed with a 15,000 resolution, 5 x $10^4$-ion target value, 100 ms maximum injection time, and 20 s dynamic exclusion. The isolation window was set to 1.6 and the normalized HCD collision energy was 38.

### 4. MS data processing

All raw data files were analyzed and processed in MaxQuant v. 1.6.17.0 (https://www.maxquant.org/) (Cox and Mann, 2008). The database search was performed with Andromeda (Cox et al., 2011) against the mouse UniProtKB database v. 2020-3 (55,462 protein entries) spiked with common contaminants and enzyme sequences. The enzyme was set as trypsin/P which cleaves after lysine or arginine even when proline follows. The search parameters included two missed cleavage sites. The variable modifications were set as methionine oxidation and protein

N-terminal acetylation. Cysteine carbamidomethylation was set as the fixed modification. The peptide mass tolerance was set to 4.5 ppm, and the MS/MS tolerance was set to 20 ppm. The false discovery rate (FDR) was set to 1% at the peptide spectrum match (PSM) and protein levels. The intensities of all ten TMT reporter ions at the MS2 level were quantified by MaxQuant using its default parameters. The TMT intensities were $\log_2$-transformed and the individual TMT channels were median-normalized. The missing values were imputed from a normal $\log_2$ reporter intensity distribution using the default settings in Perseus https://www.maxquant.org/perseus/) (width, 0.3; downshift, 1.8) (Tyanova et al., 2016). Statistical analysis of the volcano plot was performed based on the averaged $\log_2$ fold change and the Student's *t*-test p-value.
